# Microbiomes of blood feeding triatomines in the context of their predatory relatives and the environment

**DOI:** 10.1101/2023.03.24.534010

**Authors:** Hassan Tarabai, Anna Maria Floriano, Jan Zima, Natalia Filová, Joel J. Brown, Walter Roachell, Robert L. Smith, Norman L. Beatty, Kevin J. Vogel, Eva Nováková

## Abstract

The importance of gut microbiomes has become generally recognized in vector biology. This study addresses microbiome signatures in North American *Triatoma* species of public health significance (vectors of *Trypanosoma cruzi*) linked to their blood feeding strategy and the natural habitat. To place the *Triatoma* associated microbiomes within a complex evolutionary and ecological context, we sampled sympatric *Triatoma* populations, related predatory reduviids, unrelated ticks, and environmental material from vertebrate nests where these arthropods reside. Along with five *Triatoma* species, we have characterized microbiomes of five reduviids (*Stenolemoides arizonensis*, *Ploiaria hirticornis*, *Zelus longipes*, and two *Reduvius* species), a single soft tick species, *Ornithodoros turicata,* and environmental microbiomes from selected sites in Arizona, Texas, Florida and Georgia. The microbiomes of predatory reduviids lack a shared core microbiota. Like in triatomines, microbiome dissimilarities among species corelate with dominance of a single bacterial taxa. These include *Rickettsia*, *Lactobacillus, Candidatus* Midichloria, and *Zymobacter*, which are often accompanied by known symbiotic genera, i.e., *Wolbachia*, *Candidatus* Lariskella, *Asaia*, *Gilliamella*, and *Burkholderia.* We have further identified compositional convergence of analyzed microbiomes in respect to the host phylogenetic distance in both blood feeding and predatory reduviids. While microbiomes of two reduviid species from Emesinae family reflect their close relationship, the microbiomes of all *Triatoma* species repeatedly form a distinct monophyletic cluster highlighting their phylosymbiosis. Furthermore, based on environmental microbiome profiles and blood meal analysis, we propose three epidemiologically relevant and mutually interrelated bacterial sources for *Triatoma* microbiomes, i.e., host abiotic environment, host skin microbiome, and pathogens circulating in host blood.

**Importance:** This study places microbiomes of blood feeding North American *Triatoma* vectors (Reduviidae) into a broader evolutionary and ecological context provided by related predatory assassin bugs (Reduviidae), another unrelated vector species (soft tick *Ornithodor turicata*), and the environment these arthropods cohabit. For both vectors, microbiome analyses suggest three interrelated sources of bacteria, i.e., microbiome of vertebrate nests as their natural habitat, vertebrate skin microbiome, and pathobiome circulating in vertebrate blood. Despite an apparent influx of environment-associated bacteria into the arthropod microbiomes, *Triatoma* microbiomes retain their specificity, forming a distinct cluster that significantly differ from both predatory relatives and ecologically comparable ticks. Similarly, within the related predatory Reduviidae, we found the host phylogenetic distance to underlie microbiome similarities.

## Introduction

Insect species diversification has been primarily facilitated by the dietary niche expansion bound to insect physiological adaptation and the establishment of symbiotic interactions with bacteria (1, 2). Microbiomes of insects utilizing resource-limited niches (phloem, xylem, wood or blood) are known to be dominated by specialized, evolutionary old bacterial associates with biosynthetic capacities that complement the host metabolism to overcome the dietary limitations. While such symbionts may rise from taxonomically divergent groups even in closely related hosts due to frequent evolutionary replacements (3, 4), these dietary constraint have led to remarkable functional convergence. For instance, within blood sucking Anoplura, symbionts from distant bacterial genera *Lightella*, *Puchtella*, *Riesia*, *Sodalis*, *Neisseria*, and *Legionella* exhibit parallel genome evolution resulting in retention of genes coding for riboflavin biosynthesis (5, 6). Within a broader frame of blood feeding Metazoa, functional convergence was found among symbionts of unrelated insect groups, between insects and leaches (7) and were, to some extent, suggested for gut microbiomes of vampire bats and vampire finches (8).

While the majority of the approximately 7000 Reduviidae species are comprised of hemolymphophagous assassin bugs preying on other invertebrates, the subfamily Triatominae, commonly known as kissing bugs and the vectors of *Trypanosama cruzi*, converted to hematophagy on birds, mammals and reptiles (9, 10). In some respect, symbiosis of Triatominae (Reduviidae), resembles that of hematophagous vertebrates. Unlike tse-tse flies, keds, lice or bedbugs, Triatominae have not established specialized symbiosis during their evolution. They maintain relatively simple species-specific gut microbiomes that undergo remarkable ontogenetic development (11). Though there is not an ancient, universally shared microbiome among kissing bug species, kissing bugs in the genus *Rhodnius* do appear to have frequent associations with bacteria in the genus *Rhodococcus* (12), first identified as a symbiont by Wigglesworth (13). Experimental studies have provided some evidence that the association of *Rhodnius prolixus* and *Rhodococcus rhodnii* is somewhat specific as even closely related *Rhodococcus* species from other kissing bugs cannot substitute for *R. rhodnii* (14).

Both hemolymphophagy and hematophagy are highly specialized feeding strategies on prey/host body liquids that differ in composition. Invertebrate hemolymph is seemingly easily digestible, and is a rich source of free lipids and free amino acids (15). Vertebrate blood as a food source poses major challenges of heme toxicity and insufficient B vitamin content (16). Based on our knowledge from other hematophagous systems, we hypothesize that the Triatominae dietary shift from hemolymph to blood should be reflected in their microbiome structure and/or function. We further expect to reveal sources of bacterial influx associated with *Triatoma* natural habitat, i.e., host vertebrates and their nesting sites. In this study, we investigate microbiome signatures of five *Triatoma* species against the background of five species of hemolymphophagous assassin bugs. Another vector species, a soft tick *Ornithodoros turicata*, was included to the study to provide a phylogenetically independent contrast for hematophagous microbiomes sharing the same habitat. The analyzed arthropods originate from sympatric populations collected from nests of small vertebrates, primarily the white-throated wood rat *Neotoma albigula*, in the southern United States. Within this unique sample set, we track the environmental sources of the microbiomes and particularly address possible taxonomic convergence between microbiomes of sympatric populations of kissing and assassin bugs, and between the two hematophagous vectors, kissing bugs and ticks.

## Methods

### Sample set and study sites

The study was designed to encompass arthropods collected in the same microhabitat, i.e., white-throated wood rat (*Neotoma albigula*) nests, that share common phylogenetic (Reduviidae: assassin and kissing bugs) or dietary background (blood feeding vectors: soft ticks and kissing bugs). In addition, environmental samples (n=24) of the nest material were collected. Two nests (N2 and N41) housed a complete subset of the analyzed arthropod groups (Supplementary Table S1-DATASHEET S1). A total of 231 samples representing different developmental stages of kissing bugs (*Triatoma* sp., n=110), different assassin bug species (n= 37), and *Ornithodoros turicata* soft ticks (n= 84) were collected through four consecutive years (2017-2021) in the states of Arizona (n=164), Florida (n=12), Georgia (n=10) and Texas (n=45) (Supplementary DATASHEET S1). Ten additional samples of assassin bugs not associated with vertebrate nests were collected at two of the sampled localities. The localities included Las Cienegas National Conservation Area (LCNCA) and the University of Arizona Desert Station (DS) in Tucson, AZ; Gainesville, FL; Oconee National Forest (GANF) and Sapelo Island, GA; Chaparral Wild Life Management Area (Chaparral) and Lackland Air Force Base (LacklandAFB), San Antonio, TX. The sites were accessed with permissions from the relevant governing bodies (see the “Acknowledgements” section). The nest coordinates, sample type, developmental stage and sex (in adults) were recorded for acquired samples (Supplementary Table S1-DATASHEET S1).

### DNA extraction and arthropod species determination

DNA extraction of ethanol preserved samples was performed using DNeasy Blood and Tissue kit (Qiagen, Hilden, Germany) on whole abdominal tissues according to manufacturer’s instructions and stored at −75°C for the downstream molecular analysis. DNA from environmental samples was extracted with DNeasy PowerSoil Pro kit (Qiagen, Hilden, Germany). Taxonomic determination of sampled ticks was based on their indicative morphological attributes (17). *Triatoma* species were determined based on their morphological and molecular characteristics. In detail, we amplified and Sanger sequenced a 682 bp segment of the *cytB* gene with the previously published primers (18) CYTB7432F and CYTB7433R, which allowed the identification of *T. rubida*, *T. lecticularia, T. sanguisuga and T. gerstaeckeri.* Prospective T*. protracta* samples were amplified with primers TprF and TprR (11). Similarly, using previously published primers 18SF and 18SR (19), we obtained partial 18S rRNA gene sequences for assassin bugs. Details on all used primers are provided in Supplementary Table 1-DATASHEET S2. To determine assassin bug taxonomy, 18S rRNA data were aligned with MUSCLE (20) with fourteen additional sequences retrieved from NCBI (see Supplementary Table S1-DATASHEET S3 or the accession numbers). ModelTest-NG (21) was used to choose the evolutionary model for the phylogenetic inference according to Akaike’s Information Criterion (AIC) (22). The phylogenetic inference was calculated with RAxML (model GTR+I+G4, bootstrap random number seed 1234, number of bootstraps 100, random number seed for the parsimony inferences 123) (23). For the samples that were morphologically identified as the genus *Reduvius*, we employed an alternative mitochondrial marker (16S rRNA gene amplified with 16sa and 16sb primers (19)) and followed the same approach as described above.

### Amplicon library preparation

Amplification of the 16S rRNA V4-V5 region was carried out according to Earth Microbiome Project standards (EMP, http://earthmicrobiome.org/protocols-and-standards/16s/). Multiplexing of 255 samples and 15 controls utilized a double barcoding strategy including 12-bp Golay barcodes in the forward primer and 5-bp barcodes in the reverse primer. The blocking primer for 18S rRNA gene was employed as described previously (11). Seven negative controls were amplified along with the samples including two controls for the DNA extraction procedure (Blank2 and Blank21) and five controls from the PCR amplification step (NK, NK1, NK2, NK3 and NK11). Eight positive controls were used to confirm the barcoding output and to assess the detection limit and amplification bias. Positive controls included commercially purchased genomic DNA templates of four mock microbial communities with variant GC content and distribution. ATCC® MSA-1000™ (Samples MCE and MCE1) and ATCC® MSA-1001™ (Samples MCS and MCS1) are composed of 10 bacterial taxa with equal and staggered distributions, respectively. ZymoBIOMICS™ Microbial Community DNA Standard (Samples PC1 and PC3) and ZymoBIOMICS™ Microbial Community DNA Standard II (Samples PC2 and PC31) share eight bacterial and two yeast taxa with an even and log distributions, respectively. The amplicons were purified using AMPure XP (Beckman Coulter) magnetic beads and pooled equimolarly. An additional purification step using Pippin Prep (Sage science) was employed to remove high concentrations of 18S rRNA gene blocking primer and all unspecific amplification products. The nest material library containing 24 environmental templates was processed according to the same workflow with a single PCR negative control and two positive controls. The libraries were sequenced in two runs of MiSeq (Illumina) using v2 and Nano v2 chemistry with 2×250 bp output.

### Analysis of the amplicon data

Downstream processing, i.e., demultiplexing, merging, trimming, quality filtering and OTU-picking of reads, was performed by implementing corresponding scripts from USEARCH v9.2.64 as previously described (11). Briefly, merged demultiplexed reads from both sequencing runs were joined in a single dataset that was subjected to quality and primer trimming resulting in a final amplicon length of 373 bp. The OTU table was created by generating a representative set of sequences based on 100% identity clustering and performing de novo OTU picking using USEARCH global alignment at 97% identity match, including chimera removal (24). Taxonomical assignment of representative sequences was executed using BLASTn algorithm (25) against the sequences of SSU rRNA genes from SILVA_138.1_SSUREF_tax database (
https://www.arb-silva.de/no_cache/download/archive/release_138.1/Exports/) (26). Filtering of potential contaminants from the OTU table was performed using decontam package v1.18.0 (27) in R environment (28) based on frequency and prevalence methods of selection (https://rdrr.io/bioc/decontam/man/isContaminant.html) removing 48 out of 4777 OTUs (Supplementary Table S1-DATASHEETs S4 and S5). In addition, bacterial taxa of the genus *Sphingomonas* were filtered out as it was a known contaminant in our laboratory (11). A single *Wolbachia* OTU (OTU_95) determined as a contaminant by the decontam package was used for further analysis. The choice to retain *Wolbachia* OTUs in the dataset was based on their prevalence among the samples (70/232 samples) including assassin bugs (18/50) and ticks (20/50) for which associations with *Wolbachia* were previously reported (29-31).

Taking advantage of 12S rRNA data present in our dataset as a result of non-specific amplification (32), we identified blood meals from triatomines and ticks. An OTU table was generated from joined and clustered reads by utilizing USEARCH v9.2.64 (24) commands (fastq_join, fastx_uniques and cluster_otus). Taxonomic assignment of identified OTUs was performed by BLASTn search against NCBI nucleotide database (33) and restricted to the first hit only. The results were first filtered to include mammals and exclude samples with less than 20 reads. In addition, since the hits for the genera *Neotoma* and *Homo* were also observed in some negative controls, blood meal analysis was only performed with the samples where the two taxa reached higher read numbers than those in the negative controls (197 and 45 reads, respectively). The primary blood meal source was determined by the dominant 12S rRNA OTU, while the secondary sources were represented by other highly abundant OTUs found within a particular sample.

## Microbiome analyses

The profiles of the positive controls were compared and plotted using the ggplot2 v3.4.0 (34) and svglite v2.1.0 (35) packages. Additional clean up, microbiome analyses, data visualization and statistical tests were performed in R environment (28) using phyloseq v1.42.0 (36), vegan v2.6-4 (37) and MicEco v0.9.19 (https://github.com/Russel88/MicEco) and MicroEco v0.13.0 (38) packages. Graphical outputs were generated utilizing ggplot2 v3.4.0 (34) and further processed in Adobe Illustrator v25.4.1. A “clean” dataset was prepared by filtering out any archaeal, eukaryotic, mitochondrial, and chloroplast OTUs. Since the environmental samples produced considerably less amount of data, we have analyzed the “clean” dataset at two rarefaction levels (800 and 1000 reads per sample, seed=5) to retain enough of the nest material samples. We focused on effects of several microbiome determinants including environmental microbiome background of sampled habitats, host phylogenetic origin, and the dietary niche (blood versus hemolymph, and different blood meal sources).

Overall consistency of Triatominae microbiome profiles sequenced here with previously published data (11) was confirmed based on their taxonomic composition and dissimilarities found among *Triatoma* species. In MicroEco v0.13.0 (38) package, we used trans_abundance class to determine highly abundant bacterial. Alpha diversity was calculated by utilizing trans_alpha class and found differences among species were statistically evaluated based on Dunn’s Kruskal-Wallis multiple comparisons method. Using trans_beta class, beta-diversity analysis was performed based on Bray Curtis distances among individual microbiomes and visualized through non-metric dimensional scaling (NMDS) ordination. Adonis2 implemented under cal_manova method of trans_beta class was employed to confirm *Triatoma* species as a statistically significant factor driving the microbiome profile. Similarly, using trans_ubundance, trans_alpha and trans_beta classes, we have calculated alpha and beta diversity for tick, assassin bug and nest material microbiomes, including statistical evaluations for selected factors potentially underlying found variation, and we produced heatmaps to visualize their taxonomic composition.

The ps_venn function in MicEco package was utilized first to identify potentially environmentally-acquired fraction of analyzed invertebrate microbiomes (defined as OTUs present in at least 30% of all nest material samples; Supplementary Table S1-DATASHEET S6). Second, unique and shared taxa within and among different sample groups were identified under a more stringent fraction range (0.5, 0.8, 0.9 and 1 for all samples within the group). For *Triatoma* shared microbiome, the ps_venn results were visualized using the UpSet plot (https://github.com/visdesignlab/upset2 (39). The ps_venn function was further employed for identifying possible taxonomic convergence among microbiomes of blood feeding vectors (*O. turicata* and *Triatoma* spp.), related Reduviidae (kissing and assassin bugs), and within the two Reduviidae groups, i.e., among different *Triatoma* species, among different assassin species (*Reduvius sonoraensis* (“Reduvinae”), *Zelus longipes* (Harpactorinae), *Ploiaria hirticornis* (Emesinae: Leistarchini), and *Stenolemoides arizonensis* (Emesinae: Emesini).

### Phylogenetic inference of putative symbiotic taxa shared by different hosts

The analysis comprised OTUs shared between at least two different sample types (triatomines, ticks and assassins) that belong to the bacterial genera previously reported as arthropod symbionts. The datasets were generated for the families *Midichloriaceae*, *Morganellaceae*, *Rickettsiaceae* and for the genus *Asaia* from the OTU representative sequences and partial 16S rRNA gene sequences downloaded from NCBI. The datasets were aligned with MUSCLE software (20). For each alignment an evolutionary model was selected, and phylogenetic analysis performed as described above for the arthropod species determination.

## Results and Discussion

### Taxonomic determination and distribution of sampled arthropods

In total, we have molecularly identified five species of *Triatoma* spp. (*T. gerstaeckeri*, n=7; *T. lecticularia*, n=12; *T. protracta*, n=39; *T. rubida*, n=46; *T. sanguisuga*, n=6) and four species of assassin bugs (*Stenolemoides arizonensis*, n= 3; *Ploiaria hirticornis*, n=10; *Zelus longipes*, n=7; *Reduvius sonoraensis*, n=2). For 15 assassin individuals from LCNCA (AZ), 16S rRNA Sanger sequences contained signal for both the assassin and its prey, and we thus relied on morphological determination to the genus *Reduvius* (Supplementary Figure S1). Most of the analyzed *Neotoma* nests housed *Triatoma* sp. (14/16) and *O. turicata* (12/16). While ten nests allowed for comparisons between microbiomes of blood feeding vectors, i.e., ticks and triatomines, two nests served for comparisons of predatory and blood feeding Reduvidae. Another two nests contained some species from all three sample types, i.e., ticks, assassins and triatomines. In addition, *Zelus longipes* (Harpactorinae) and *Ploiaria hirticornis* (Emesinae: Leistarchini) were found in two locations in Georgia not associated with *Neotoma* nests (Supplementary Table S1-DATASHEET S1).

### Quality of rRNA amplicon data

The MiSeq runs resulted in 5,998,041 sequence pairs with successful merging and trimming of 3,278,636 pairs (54.66%) sharing a mean merged length of 404-bp and an average number 7514 reads per sample. The profiles of negative controls were utilized to identify contaminant OTUs (see Materials and Methods, Supplementary Table S1-DATASHEET S4). The sequencing of the positive controls showed a bias against *Rhodobacter* and in favor of *Staphylococcus* species, but the overall frequencies of the sequenced taxa are consistent with their abundance in the input material and comparable across the control replicates (Supplementary Figure S2). OTUs (n=4,777) were clustered from the entire dataset, i.e., amplicons retrieved for 255 arthropods, related environmental material and controls (Supplementary Table S1-DATASHEET S5). For 12S rRNA gene, used for determination of primary and secondary blood meal source (Supplementary Table S1-DATASHEET S7), a total of 1,287,068 reads were retrieved within the arthropod dataset with an average of 5,297 reads per sample.

### Microbiomes of North American *Triatoma* kissing bugs

The profiles of *Triatoma* microbiome analyzed here concur with the characteristics previously published by our team (11). These include significant differences in taxonomic composition and diversity among different *Triatoma* species (Supplementary Table S1-DATASHEETS S8-S10) which undergo an ontogenetic shift from a diverse microbiome to a very simple community dominated by a single bacterial genus (Supplementary Figure S3). Venn analyses performed here confirmed Actinobacteria as the dominant phyla in *Triatoma* microbiomes with the genus *Dietzia* found in over 50% individuals of phylogenetically closely related *T*. *rubida*, *T*. *lecticularia*, and *T*. *protracta.* The same fraction of *T*. *gerstaeckeri* and *T*. *sanguisuga* individuals shared two actinobacterial genera, i.e., *Pseudonocardia* and *Nocardioides* (Supplementary Table S1-DATASHEET S11).

Regardless of the *Triatoma* phylogenetic relationships, *T*. *gerstaeckeri*, *T*. *lecticularia*, *T*. *rubida* and *T*. *sanguisuga* shared two proteobacterial taxa, an *Acinetobacter* (OTU_17), and *Serratia* sp. (OTU_14, Supplementary Table S1-DATASHEET S11). While we cannot currently assess their role in *Triatoma* microbiomes, it is worth noting that both taxa are commonly found in insects (40, 41) and in some cases profound advantages have been described. For instance, some *Acinetobacter* strains are associated with insect resistance to the insecticide cypermethrin (42), a pyrethroid used for control of insect pests (43-45). *Serratia marcescens* isolated from Triatominae microbiome poses negative effects on survival and replication of *Trypanosoma cruzi* in *Rhodnius prolixus* gut (46, 47). Notably, number of other bacterial taxa are almost universally present in *Triatoma* species across the majority of individuals (≥ 80%), for instance *Staphylococcus, Massilia, Bacillus, Planoccoccus, Streptomyces* (Figure 1). Most were previously reported in microbiomes of some Triatominae species (11, 48, 49). We elucidate putative origin of these omnipresent taxa in the following sections, providing evolutionary and ecological frame of this study.

**Figure 1:**
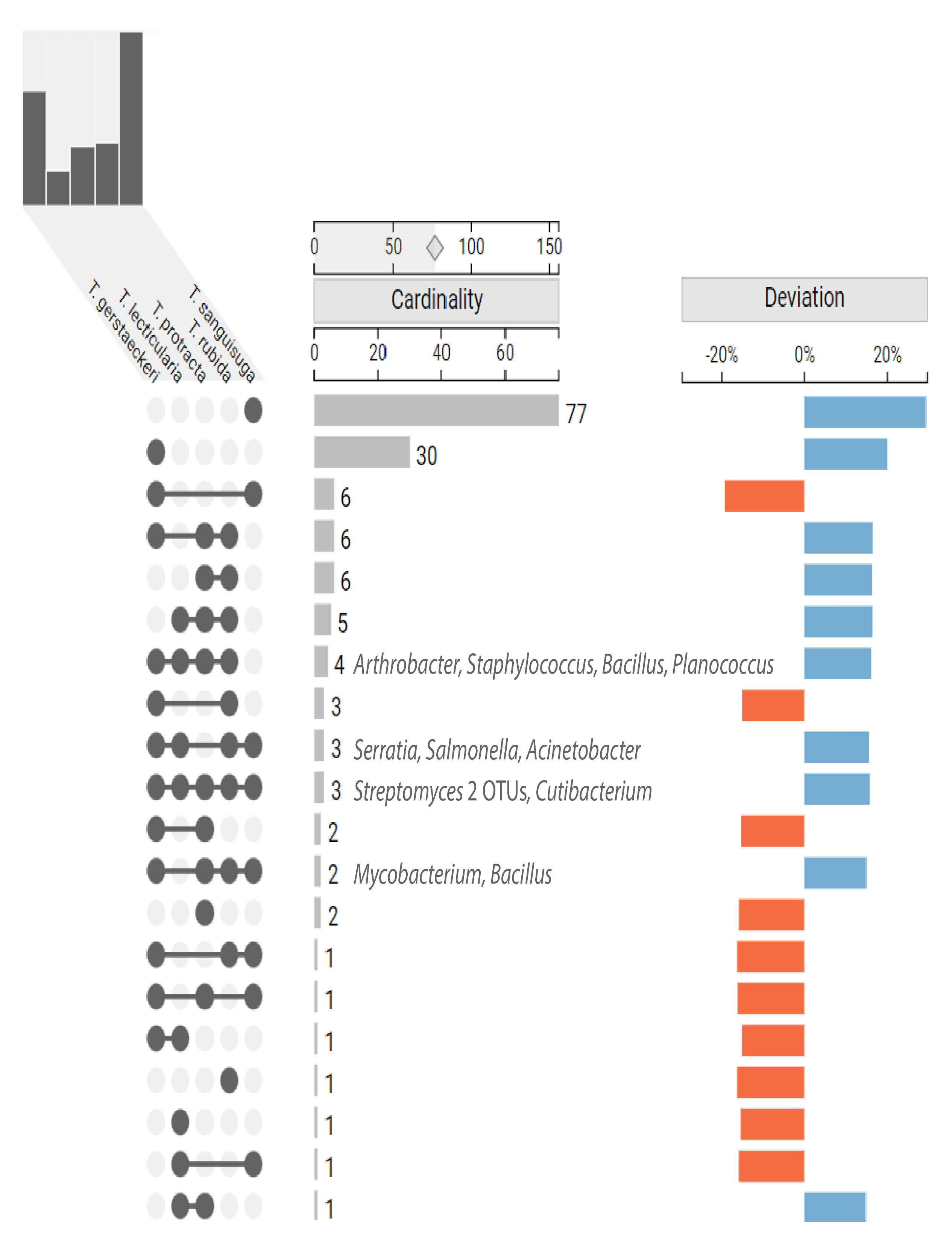
UpSet graph depicting microbiome intersections among different *Triatoma* species. The top bar chart stands for the cumulative number of OTUs identified for each *Triatoma* species. The colored plot indicates how much each intersection deviates from the expected size if species membership within the analyzed dataset was random. The bacterial genera shared, in 0.5 fraction, by four and more species are listed.

### Microbiomes of related predatory assassin bugs

Similarly to *Triatoma*, host species was a significant predictor of assassin bug microbiome alpha (Supplementary Table S1-DATASHEET S12) and beta diversity (Figure 2, Supplementary Table S1-DATASHEETs S13 and S14) explaining over 60% of variation. Clear dissimilarities corelate with dominance of a single bacterial taxa in each assassin bug species, i.e., *Rickettsia* in *Ploiaria hirticornis*, *Lactobacillus* in *Reduvius* sp.*, Candidatus* Midichloria in *Stenolemoides arizonensis*, and *Zymobacter* in *Zelus longipes* (Figure 2A). Species specificity and a single taxon dominance resemble the microbiome characteristics of related hematophagous kissing bugs (11) but pose a sharp contrast to the recently published microbiome patterns in Old World assassins (50). Based on the data from six Harpactorini species, Li and colleagues suggested *Enterococcus* bacteria to be a conserved microbiome component to most assassin bugs (50). In our data, *Enterococcus* is however only found in some individuals of *Reduvius* sp. and *Stenolemoides arizonensis,* both distantly related to the Harpactorini group of reduviids. For the only Harpactorini species in our dataset, *Zelus longipes*, *Enterococcus* presence is revoked across all individuals. The same applies for analyzed *Ploiaria hirticornis* individuals from Emesinae group. A surprisingly high number of assassin bug-associated bacteria may be of further interest as they belong to known insect symbionts. These are unevenly distributed and include *Rickettsia, Candidatus* Midichloria, *Wolbachia*, *Candidatus* Lariskella, *Asaia*, *Gilliamella*, *Burkholderia*, and the bacteria from *Morganellaceae* family (designated as “endosymbiont8”; Figure 2A) with the closest BLASTn hit to *Arsenophonus* symbionts known for a broad range of associations with invertebrates (51).

**Figure 2:**
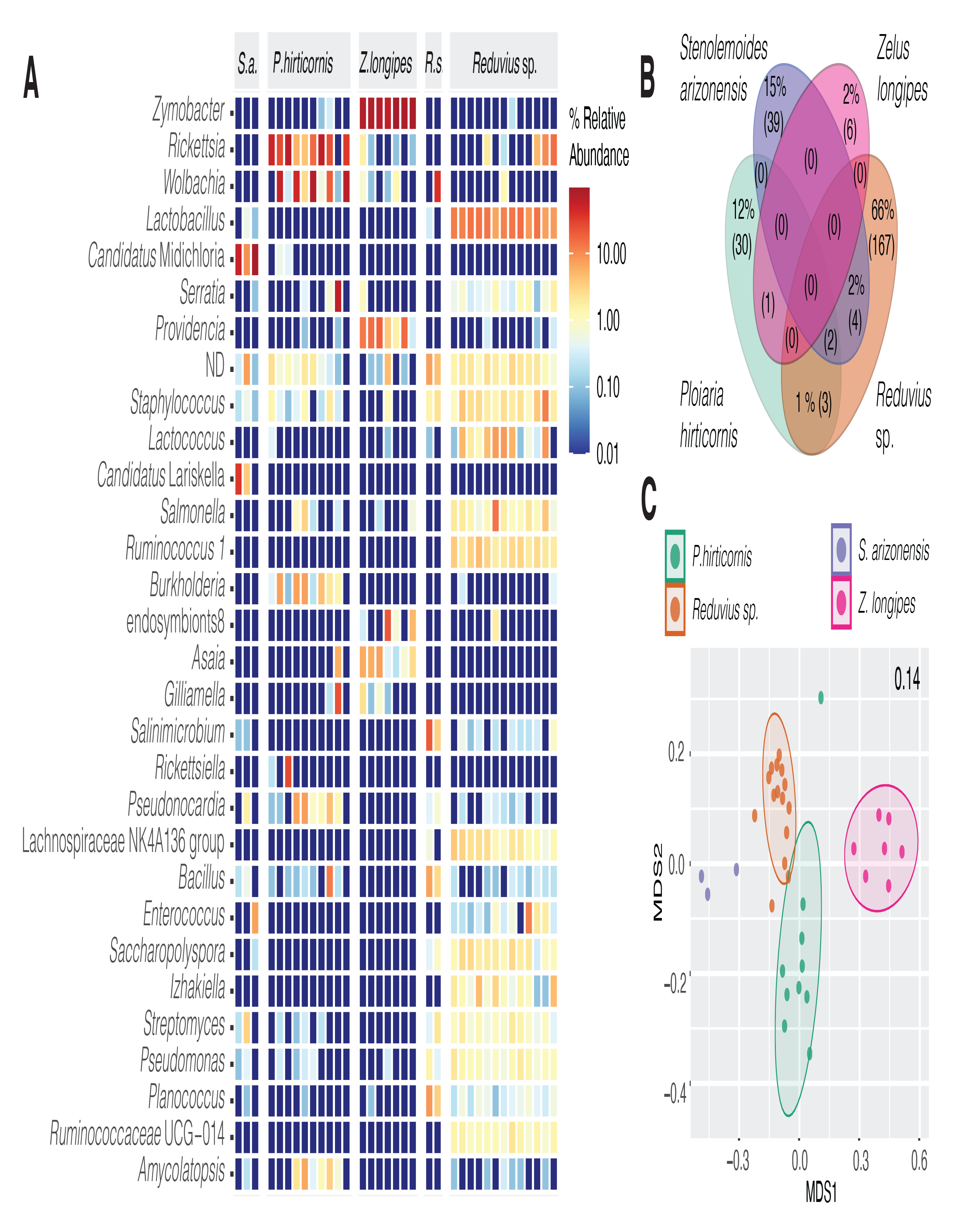
Microbiome profiles of hemolymphophagous assassin bugs. Heatmap showing the distribution of the 30 most abundant bacterial taxa identified across the assassin species (**A**). Shared proportion of the microbiome components among the analyzed assassins. The numbers in the parentheses stand for the absolute number of OTUs. Shared proportion of the microbiome that accounts for less than 1% is only depicted as the absolute count (**B**). NMDS clustering based on Bray-Curtis distances. The number in the right upper corner indicates the stress value. (**C**). Two *Reduvius* species were merged under a single group, *Reduvius* sp.

However, the primary goal of this study was not a thorough analysis of microbiome components in predatory Reduviidae. We are only providing a snapshot from the vast diversity of known reduviids that serves as habitat specific frame for investigation on microbiome signatures underlined by evolutionary significant dietary switch of Triatominae predecessors. While the shared component analysis among surveyed assassins does not suggest existence of assassin core microbiome (Venn for 0.5 fraction; Figure 2B, Supplementary TableS1-DATASHEET S15), under less stringent conditions we have identified the genera *Salmonella* and *Serratia* as shared among *Triatoma* and some assassins (Supplementary Table S1-DATASHEET S16).

### Microbiome of the soft tick *Ornithodoros turicata*

The microbiome profiles retrieved in this study resemble of those described for *O. turicata* from Bolson tortoises in northern Mexico (52). When compared among the three sampled localities, microbiomes of Lackland collected ticks displayed significantly higher measures for alpha diversity (Figure 3C, Supplementary Table S1-DATASHEET S17). In the ordination-based analysis, the microbiomes clustered according to their geographic origin (PCoA capturing 16% of the variation, Figure 3B, Supplementary Table S1-DATASHEET S18 provides statistical support). Compared to the results of Barraza-Guerrero and colleagues, *Midichloria* related symbionts were not the most abundant taxa in our dataset. Several individuals however harbored two *Candidatus* taxa from Midichloriaceae family, *Jidaibacter* and *Lariskella* (Figure 3A). Here, microbiomes were dominated by the known symbiotic genera of ticks, i.e., *Coxiella* and *Rickettsia* (Figure 3). While *Coxiella* symbionts have been previously described from *O. turicata* and other *Ornithodoros* species (52, 53), bacteria of the genus *Rickettsia* were only identified as potential symbionts of another soft tick species, *Carios vespertilionis* (53). The distribution of these two genera across our dataset differed substantially. The genus *Coxiella* was omnipresent across nests and localities, being detected in 96% of analyzed individuals. The genus *Rickettsia* was only associated with 26 individuals sampled in Desert Station, AZ (32% of all the samples). *Rickettsia* presence/absence further reflects the origin of the analyzed ticks from individual *Neotoma* nests and suggests the bacteria may rather represent vertebrate pathogens acquired through feeding. A similar pattern was observed for *Borrelia* pathogens vectored by ticks. *Borrelia* nest specific distribution was further corroborated by presence of the pathogen (detectable at median relative abundance of 0.27%) in *Triatoma* individuals cohabiting the same nests as the infected ticks.

**Figure 3.**
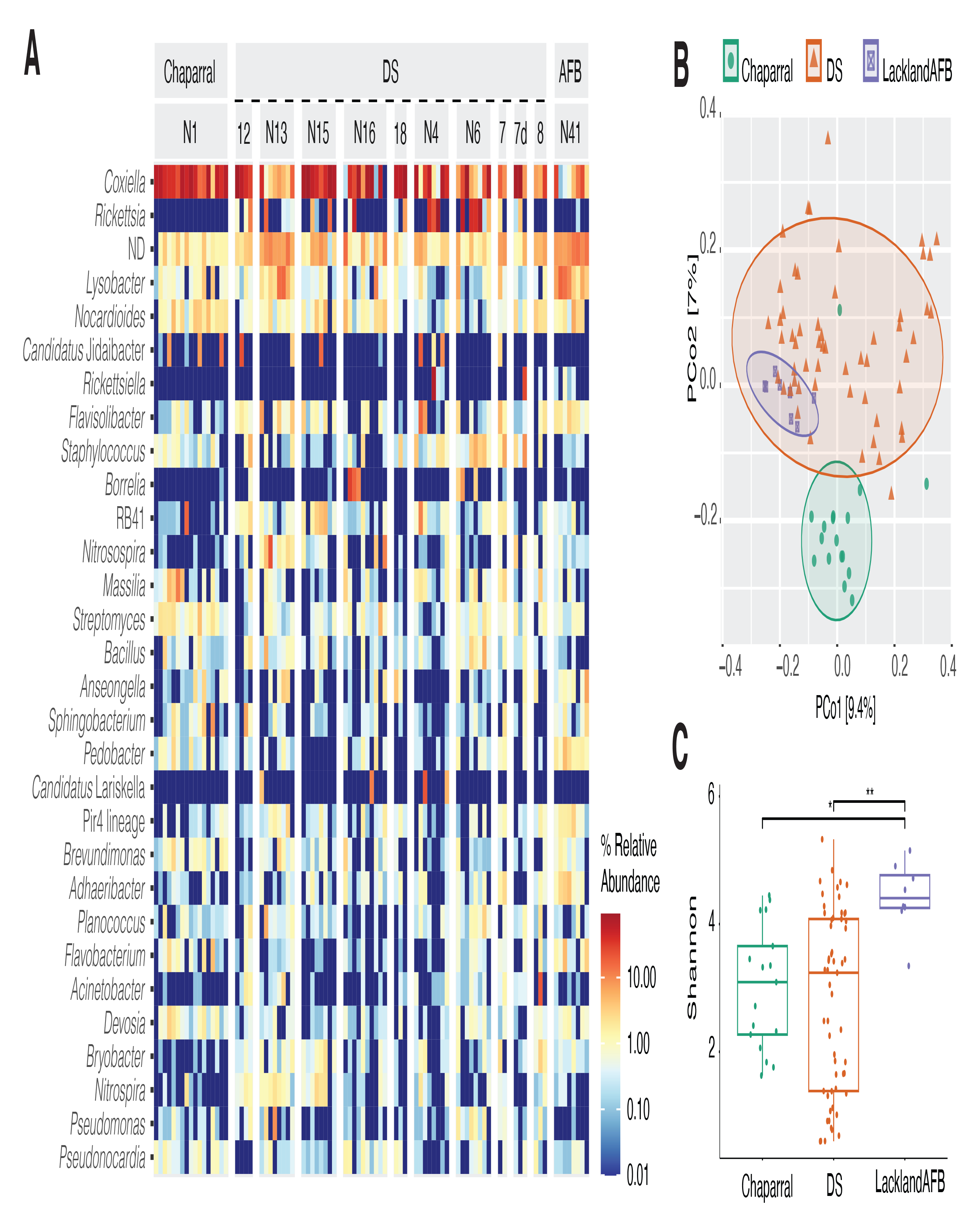
*Ornithodoros turicata* microbiome profile. Heatmap for the 30 most abundant genera, ND stands for sum of the unclassified taxa at the genus level (**A**). Principal coordinates analysis based on Bray Curtis distances, colored shapes stand for different localities: DS- Desert Station, Tucson, AZ; Chaparral-Chaparral WLMA, TX; AFB-Lackland Air Force Base, San Antonio, TX (**B**). Bray Curtis microbiome dissimilarities calculated among the three localities (**C**). Asterisks indicate significant differences found between the diversity measures (Dunn’s Kruskal-Wallis Multiple Comparisons, Supplementary Table S1-DATASHEET S17).

### Phylogenetic inference of possibly symbiotic taxa shared by different hosts

We investigated phylogenetic origin of possibly symbiotic OTUs from the families *Candidatus* Midichloriaceae, *Morganellaceae*, *Rickettsiaceae* and *Acetobacteraceae* detected across different hosts. For the *Rickettsiaceae* OTUs, present in all *P. hirticornis* and *Reduvius* samples, five *Zelus* samples, and 37 *O. turicata*, our amplicon data could not provide sufficient phylogenetic resolution (Supplementary Figure S5). Similarly uncertain result was obtained for *Acetobacterceae* OTUs taxonomically assigned to the genus *Asaia*. While OTU_39 falls among symbionts of mosquitos and *Asaia* species isolated from flowers (54, 55) the relationship of two other sequences to the genus remains highly questionable (Supplementary Figure S6).

Three *Midichloriaceae* OTUs, found in higher abundances in 14.2% (n= 5/35) of the assassin bugs, 20.7% (n=17/82) of ticks, correspond to three distinct lineages (Figure 4), two of which encompass endosymbionts of arthropods. OTU_6, present in assassin bugs, falls within *Midichloria* genus, a well-known group of tick endosymbionts (56). OTU_26 found in both assassin bugs and ticks clusters with the *Lariskella* group, which includes symbionts of ticks and insects (57). Interestingly, OTU_16 present in ticks clusters with two *Fokinia* species, which reside in aquatic environment as symbionts of *Paramecium* (58, 59). However, this node is not highly supported, including two highly diverging *Midichloriaceae* (58, 59), and might be a product of long branch attraction. For the *Morganellaceae* OTU we prepared a dataset of 154 16S rRNA gene sequences, as in (51), including a single outgroup. The inferred phylogeny (Supplementary Figure S7) is well supported and clearly shows OTU_67 clustering with *Morganellaceae* symbionts of aphids. Among *Zelus longipes* individuals the OTU distribution does not however suggest that the bacteria play a stable symbiotic role (being detected in 50% of the samples) and may point out its origin in the prey that often includes aphids.

**Figure 4:**
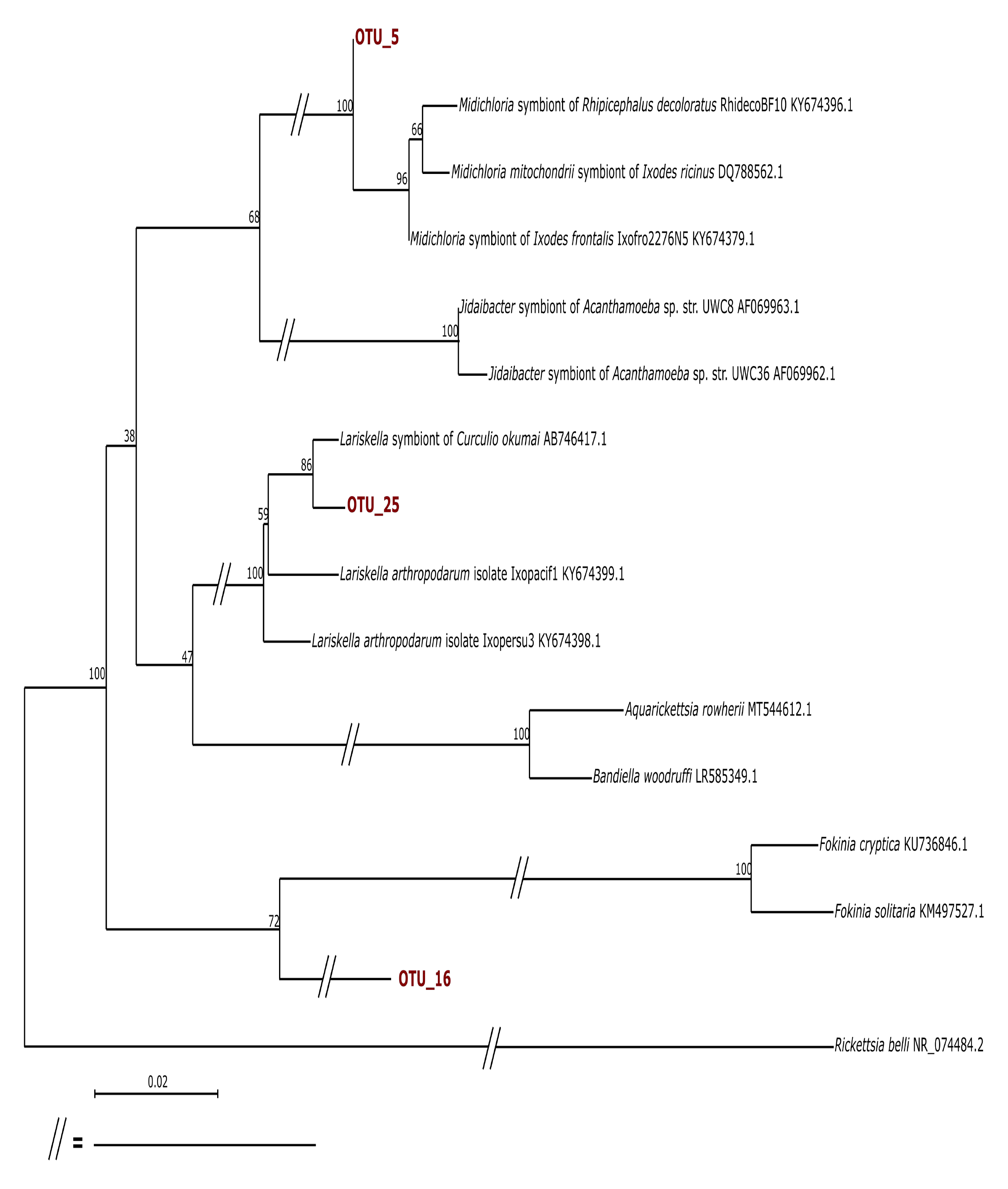
Phylogenetic inference on 16S rRNA gene sequences of *Midichloriaceae* OTUs associated with assassin bugs and ticks. Numbers at the nodes are bootstrap values.

### Blood meal signature among *Triatoma* and tick microbiome

Primary blood meal was identified in all five sampled *Triatoma* species (93 individuals, 84%), and five individuals (6%) of *Ornithodoros turicata* (Supplementary Table S1-DATASHEET S7). The striking difference in the fractions with determined vertebrate host is undoubtedly related to the distinct physiology of triatomines and soft ticks. While both vectors feed on similar hosts, including mammals, reptiles and birds (10, 60), soft ticks are known to endure extremely long starvation periods (61) that may hamper the molecular detection of ingested blood meal (62). For *T. protracta* (38/39), *T. rubida* (34/46) and *T. lecticularia* (10/12), packrats, *Neotoma* sp., were indeed the most common blood meal (82/93, 88%). For six triatomines in our dataset (6%), i.e., *T. gerstaeckeri* (4/7) *and T. sanguisuga* (2/6), armadillos, *Dasypus* sp., served as a blood meal source. Five *Triatoma* individuals fed on species of brown-toothed shrews, small insectivores in the genus *Episoriculus sp.* (3), and rodents of the genera *Peromyscus* (deermice) and *Callospermophilus* (ground squirrels). For 35 *Triatoma* individuals and a single tick, we were able to determine secondary blood meal sources (see Materials and Methods). These included the genera *Felis* (n=32), *Dasypus* (n=3), and individual records on *Callospermophilus* and *Neotoma* (Supplementary Table S1-DATASHEET S7). *Neotoma* predominance along with the absence of human and domestic animals, may well reflect the sampling strategy, i.e., active search for triatomines in sylvatic packrat nests. The range of other less frequent hosts listed above is in line with earlier studies on North American *Triatoma* species (63, 64).

The uneven distribution among identified blood sources did not allow us to further evaluate whether different vertebrate hosts may determine the vector microbiome profile, as previously discussed for triatomines (65, 66) and other vectors, e.g., yellow fever mosquito (67), tsetse flies (67), and western black-legged ticks (68). We therefore searched for common microbiome signatures among those individuals that fed on *Neotoma* (Supplementary Table S1-DATASHEET S19). At least 50% of *Neotoma* fed *Triatoma* and tick individuals housed *Cutibacterium* (OTU_41), *Streptomyces* (OTU_102 and OTU_295), *Brachybacterium* (OTU_650), *Arthrobacter* (OTU_214), *Microbacterium* (OTU_254), *Pseudonocardia* (OTU_513), *Massilia* (OTU_43), *Pseudomonas* (OTU_59) and *Acinetobacter* (OTU_17) pointing out their potential link to the vertebrate host. Three of the OTUs belongs into antimicrobial-producing bacteria (i.e., Pseudonocardiaceae and Streptomycetaceae) previously identified within Key Largo woodrat (*Neotoma floridana smalli)* body microbiome and the microbiome of their nests in Florida (69, 70). In addition, *Cutibacterium*, a common skin commensal from Propionibacteriaceae family (71), has been repeatedly recorded in the body swabs from small rodents (unpublished laboratory data). Notably, all six *Triatoma* individuals that fed on *Dasypus* harbored *Alteribacillus* (OTU_118), *Pseudonocardia* (OTU_177) and *Mycobacterium* (OTU_2341). The *Mycobacterium* OTU identified here represents a good candidate for blood meal related microbiome signature since *Mycobacterium leprae*, the causative agent of leprosy in humans, is commonly found among nine-banded armadillo (*Dasypus novemcinctus*) in Texas and Florida (72).

### Environmental microbiome

The evaluation of nest material revealed significant amount of variation between microbiome diversity of nests sampled in Gainesville, FL and Lackland AFB, TX (adonis2: R^2^= 0.274, P=0.008; Supplementary Figure S4, Supplementary Table S1-DATASHEETS S20 and S21). The location also determined significant differences found for all alpha diversity measures (Dunn’s Kruskal-Wallis Multiple Comparisons, Supplementary Table S1-DATASHEET S22 and Supplementary Figure S4). While the nest microbiome from Florida tends to be dominated by a few taxa, especially Bacilli genera *Peanibacillus*, *Bacillus* and *Lysinibacillus*, Texas locations display more diverse and equally structured environmental microbiome (Figure 5A) composed mainly of Acidobacteriia (*Bryobacter*, *Edaphobacter*), Chitinophagia (*Ferruginibacter*, *Flavisolibacter*), and Planctomycetia (*Singulisphaera* and *Pirellula* related Pyr4 lineage). Few taxa are, in different relative abundances, universally present across the sampled nests, i.e., *Peanibacillus, Bacillus,* and *Planococcus* (Figure 5A). As the universal components of nest microbiomes, *Bacillus* (OTU_20) and *Planococcus* (OTU_50) are, at different Venn fractions, also observed in samples of *Triatoma*, ticks and assassin bugs (Figure 5B, Supplementary Table S1-DATASHEET S23 and S11), supporting a role of environment in shaping the *Triatoma* microbiomes as suggested previously (11). For triatomines and ticks occupying the same nest, *Massilia* (OTU_43) and several *Streptomyces* OTUs (Figure 5B) were found as shared microbiome components, corroborating the findings on *Neotoma* fed vectors above. Two OTUs, i.e., *Staphylococcus* (OTU_3) and *Cutibacterium* (OUT_41) were found as common across different arthropods from different nests (Figure 5B).

**Figure 5:**
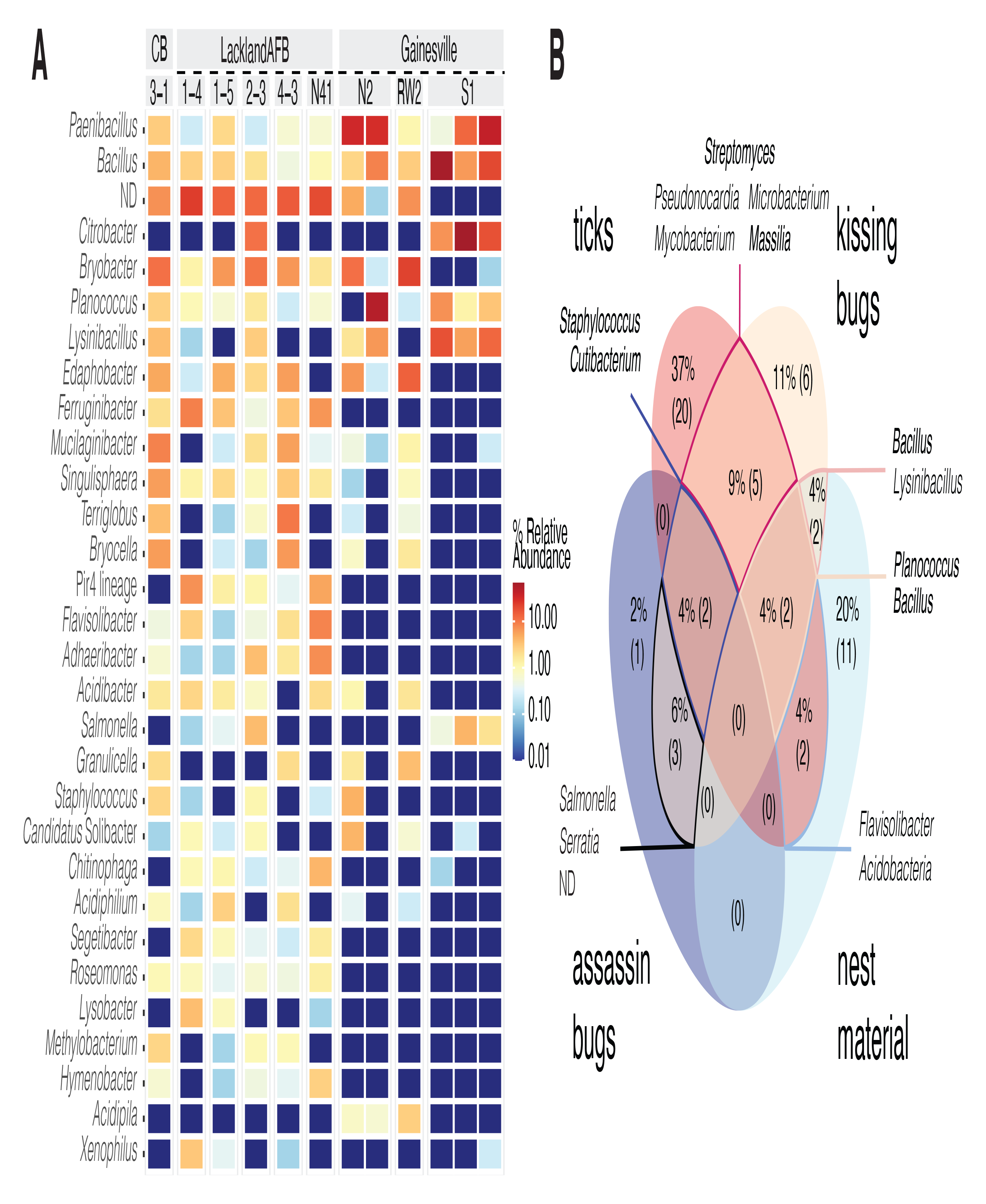
Nest microbiome characteristics. Heatmap for the 30 most abundant genera across the sampled nests, ND stands for sum of the unclassified taxa at the genus level (**A**). Venn analysis at 0.5 fraction among all sample types, i.e., ticks, kissing and assassin bugs, and nest material (**B**). The numbers in the parentheses stand for the absolute number of OTUs. The taxa in bold, were repeatedly recognized as shared microbiome members under different fractions (Supplementary Table S1-DATASHEETS S22 and S10) and thus considered of nest environmental origin.

Since our nest material sampling does not fully mirror the arthropod sample set, especially lacking nest data from Arizona, we are limited in differentiating the environmental component within all analyzed arthropod microbiomes. In triatomines and ticks, the above-mentioned taxa, commonly found in skin or soil microbiomes (73-76), likely originate from host-nest ecological interface. Based on the nest material and blood meal analyses we thus suggest three mutually interrelated bacterial sources for vector microbiomes (Figure 6). First, environmental microbiome of their natural habitat, i.e., vertebrate nests and middens, as suggested previously in *Triatoma* (11) and other true bugs (50). Second, vertebrate skin and skin-derivate microbiomes (77-79) that vectors encounter while feeding (*Neotoma* Supplementary table S1-DATASHEET S19). Third, vertebrate pathogens circulating in blood as illustrated here for *Triatoma* on *Mycobacterium – Dasypus* blood meal link, and also supported by the distribution patterns for *Rickettsia* and *Borrelia* OTUs across the unfed ticks (see Microbiome of blood feeding vector *Ornithodoros turicata*). While in *Triatoma* and *O. turicata* microbiomes these bacteria likely represent transitional components, in other hematophagous vectors, like ticks and lice, vertebrate pathogens gave rise to evolutionary stable symbiotic associations (80-82).

**Figure 6:**
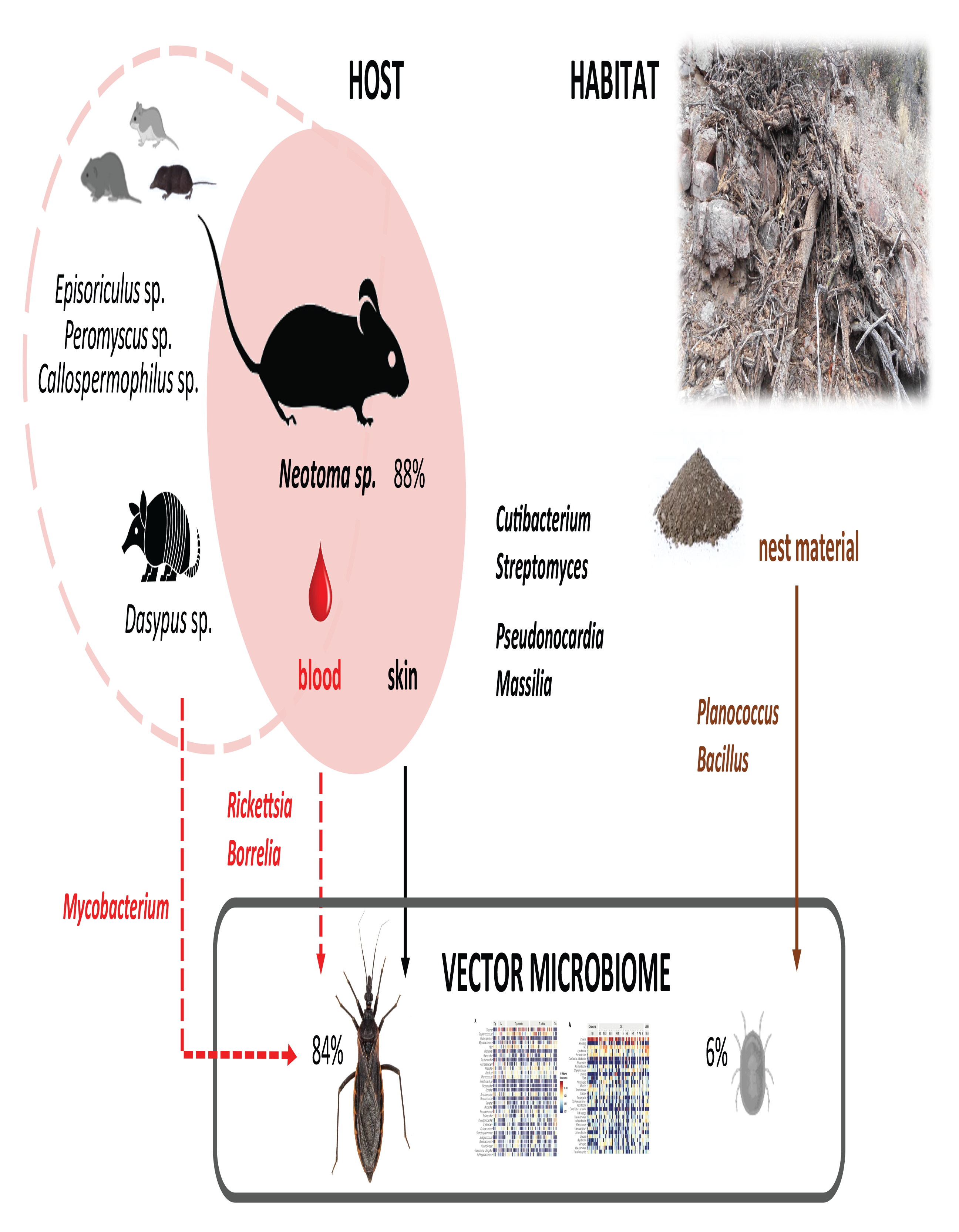
Scheme for putative sources of vector microbiome based on bloodmeal analysis and nest microbiome profiling. Percentages next to each vector stand for successfully identified primary blood meal from which 88% matched *Neotama* sp. The four taxa (*Cutibacterium, Streptomyces, Pseudonocardia* and *Massilia*) might originate from both the host skin and the nest microhabitat.

### Host phylogeny and diet as the microbiome determinants

Clustering among analyzed arthropod microbiomes based on their similarities (calculated as weighted Unifrac distances, Figure 7) shows triatomines in a monophyletic cluster and a close relationship between the microbiomes of two Emesinae species. These data suggest compositional convergence of microbiomes in respect to the host phylogenetic distance. The phylogenetic position of *O. turicata* soft ticks as a distant outgroup for the analyzed true bugs is not reflected in the microbiome dissimilarities. *O. turicata* clusters within the hemolymphophagous assassin bugs, resembling the Emesinae microbiomes dominated by common members of Rickettsiales. While we have identified a shared microbiome component among the two hematophagous vectors (see Blood meal signature among *Triatoma* and tick microbiomes), it encompasses mostly less abundant taxa of putatively environmental origin or blood born pathogens. These patterns are well reflected by clustering in the NMDS analysis and statistical tests supporting the host specificity of *Triatoma* microbiomes. The sample type acts as a significant predictor of microbiome profile, explaining close to 26% of found variation (Supplementary Table S1-DATASHEET S24). Diet was identified as another significant predictor, though the effect is comparably lower accounting only for 6% of found microbiome variation (Supplementary Table S1-DATASHEET S24). Both diet and phylogenetic distance as factors underlying Triatominae microbiome structure indirectly support our hypothesis on microbiome change bound to Triatominae evolutionary shift from hemolymphophagy of predatory Reduviidae to blood feeding. Since the methodology employed here limited our focus on the microbiome composition and structure, functional convergence between microbiomes of blood feeding and other predatory Reduviidae remains to be evaluated with complex metagenomic approaches.

**Figure 7:**
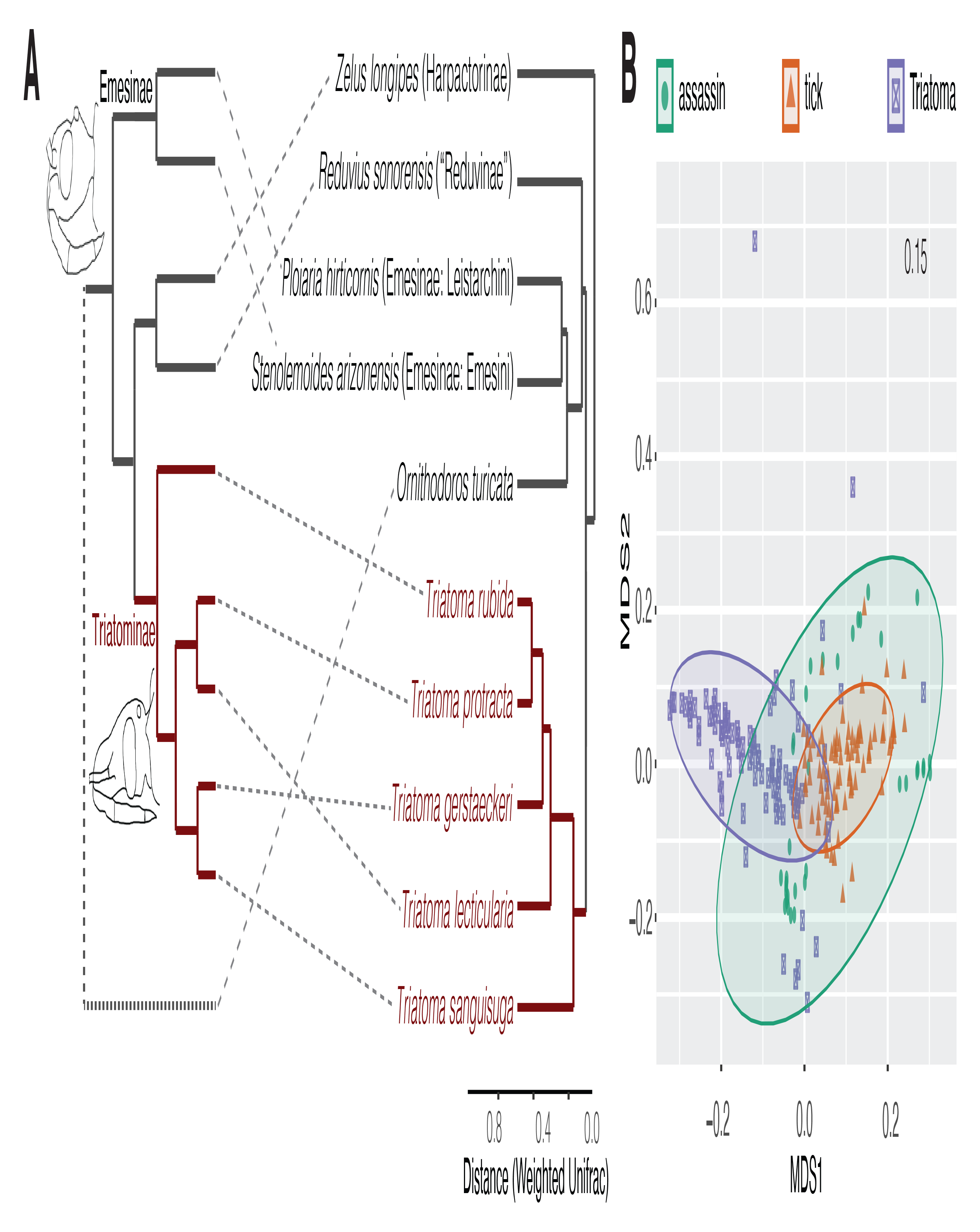
Schematic host phylogenetic relationships compared to the microbiome dissimilarities based on Weighted Unifrac distances (**A**). The individual microbiome distances are visualized using NMDS (**B**).

## Supporting information

Supplementary Table S1-DATASHEETs S1 to S24

## Acknowledgement

We would like to acknowledge the University of Arizona, Department of Entomology’s Desert Station for graciously supporting our field activities. We thank Samantha M. Wisely and Chanakya R. Bhosale for logistical and technical assistance with sampling in Gainesville, FL. We further acknowledge the contributions of the Texas Park and Wildlife Management Department and the Bureau of Land Management (Gila District Office, Tucson, AZ). This work was supported by the Czech Science Foundation (grant number 21-10185M to E.N.).

## Contributions

EN conceived the study. EN, AMF, JJB, NF, RLS, NLB, KV and WR participated in planning and executing field collections. JZ and NF performed the DNA isolation, library preparation and sequencing. HT, AMF and EN analysed the data and interpreted results. HT, AMF and EN wrote the draft and all the authors contributed to its improving. All the authors read and approved the final manuscript.

## Competing interests

All the authors declare that they have no competing interests. Robert L. Smith has not sought nor has he received any remuneration for providing access to Desert Station and adjacent private land and residence, nor has he received payments from any researchers or scholarly visitors on Desert Station.

## Data Availability

Raw sequence reads generated from this study were deposited in the NCBI Sequence Read Archive (SRA) repository under Bioproject PRJNA898622. The complete R code employed in this study with its associated datasets are available at https://github.com/hassantarabai/Convergency-MS-2022.

## Notes

### Competing Interest Statement

The authors have declared no competing interest.

