## Supplementary Figures for "Microbiomes of blood feeding triatomines in the context of their predatory relatives and the environment"

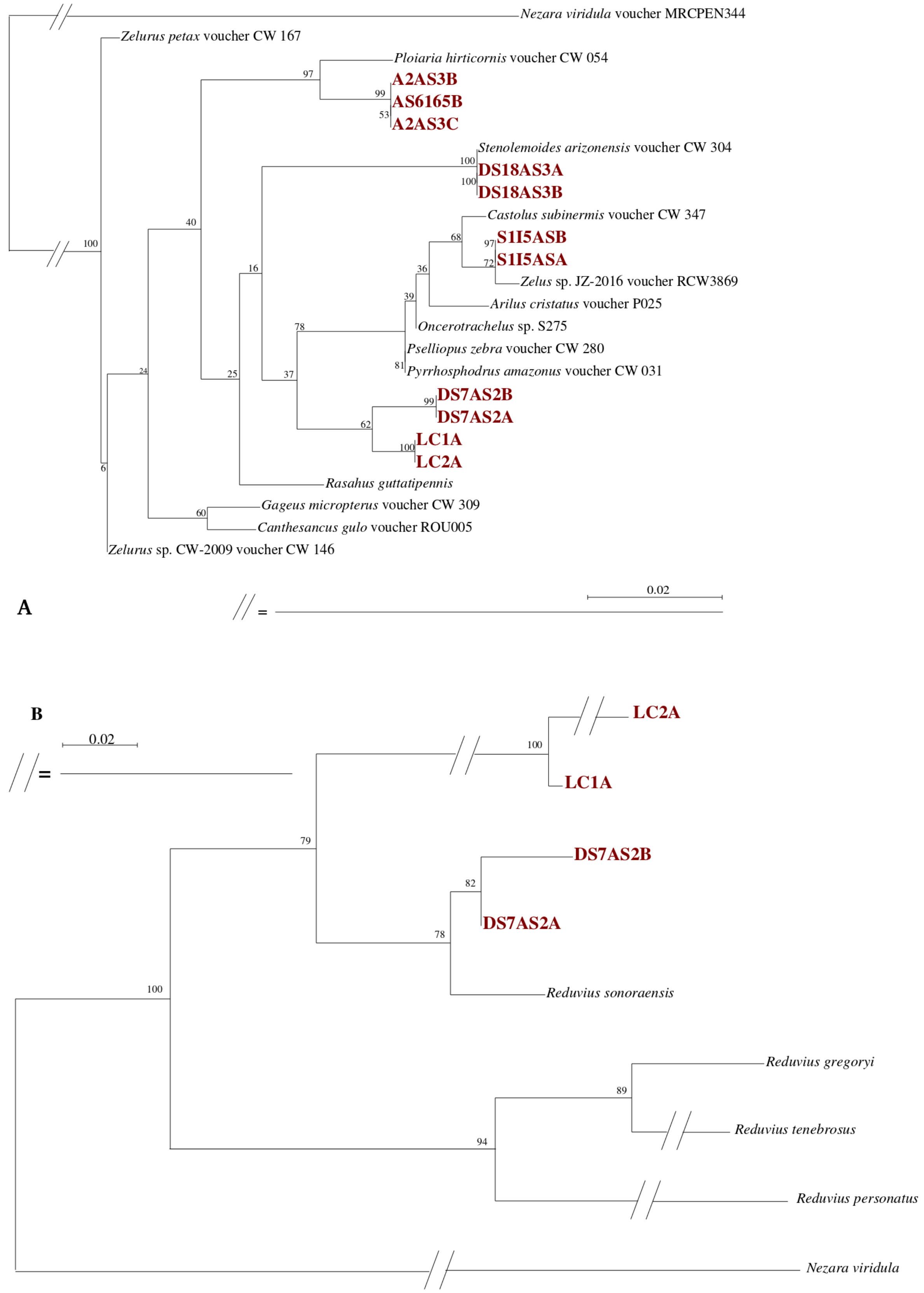


**Supplementary Figure S1:** Taxonomy assignment of sampled assassin bugs. Phylogeny inference on 18S rRNA gene sequences (A) did not allow for complete species determination. 16S rRNA gene-based inference was performed for the remaining four samples (B).

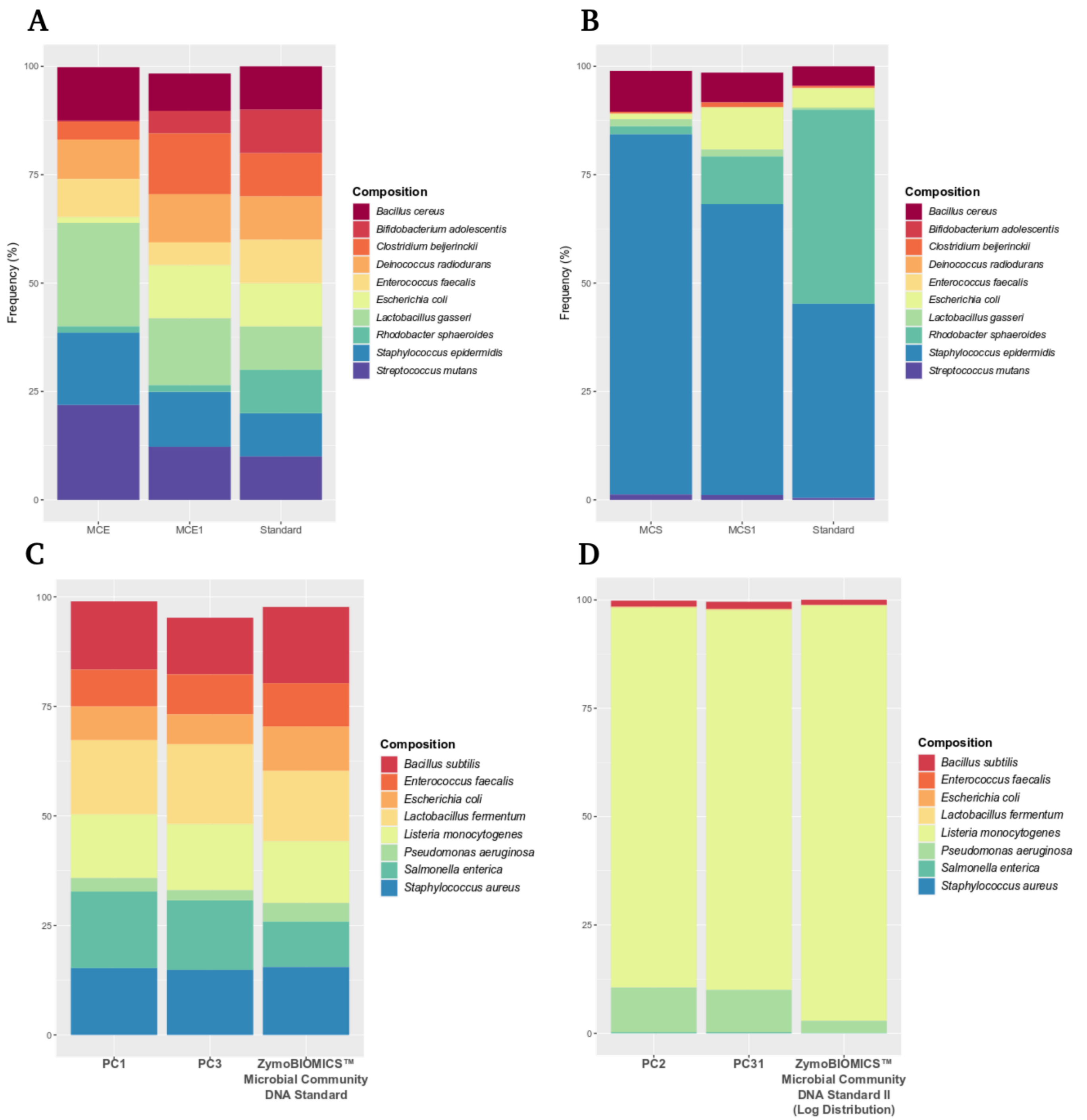


**Supplementary Figure S2:** Frequencies of taxa in the positive controls as recovered by 16S rRNA amplicon sequencing. In every plot, columns on the right represent the frequencies declared in the commercially purchased DNA templates. **A:** ATCC® MSA-1000™ Standard, **B:** ATCC® MSA-1001™, **C:** ZymoBIOMICS™ Microbial Community DNA Standard II, **D:** ZymoBIOMICS™ Microbial Community DNA Standard II Log Distribution. Overall, frequencies are coherent, albeit under-sequencing of *Rhodobacter sphaeroides* and over-sequencing of *Staphylococcus epidermidis* and *Streptococcus mutans* (A, B), and over-sequencing of *Pseudomonas aeruginosa* (D).

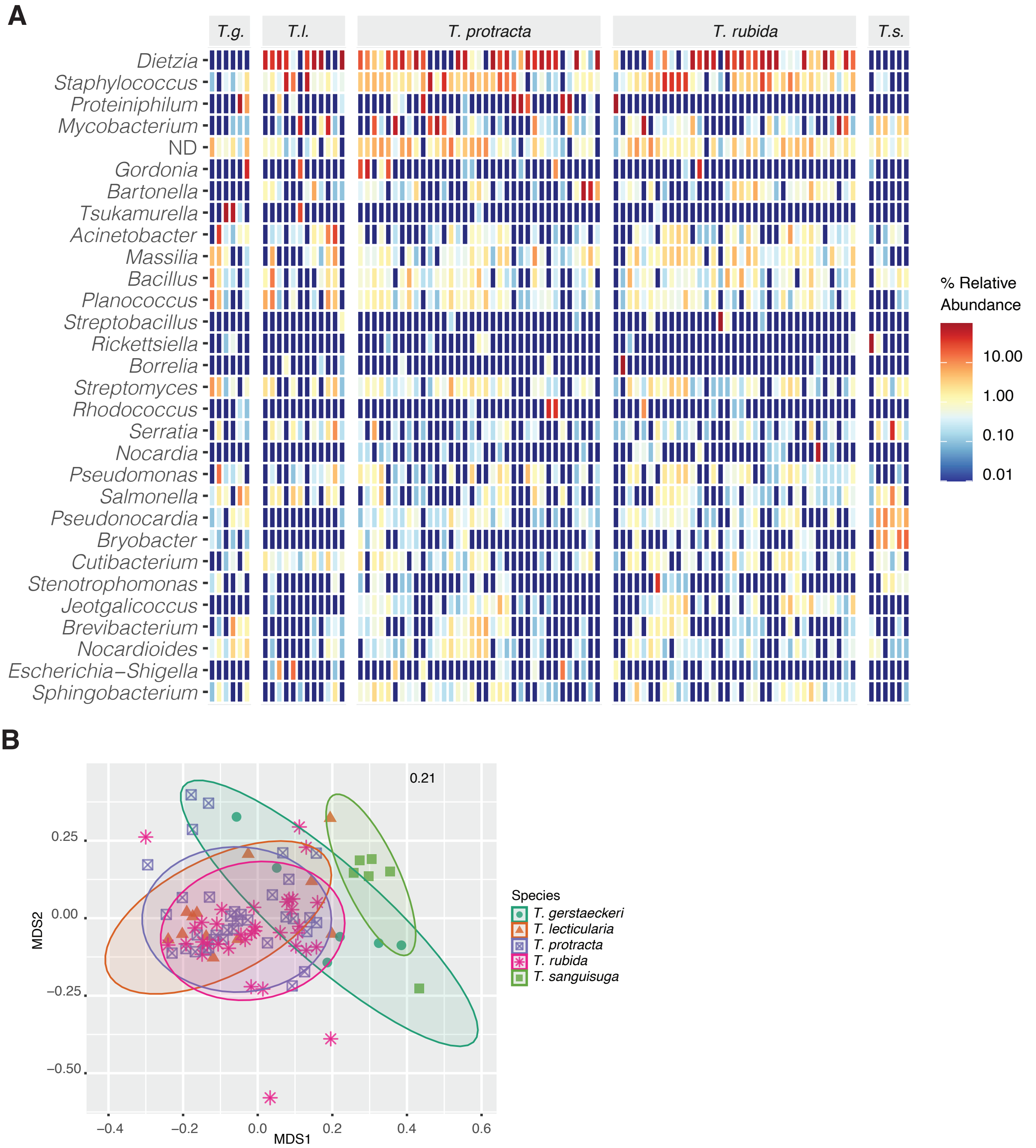


**Supplementary Figure S3:** Microbial profiles of five *Triatoma* species. **A:** Heat map for first 30 most abundant genera across the *Triatoma* samples. **B:** NMDS ordination for based on Bray-Curtis dissimilarities. The number in the right upper corner indicates the stress value.

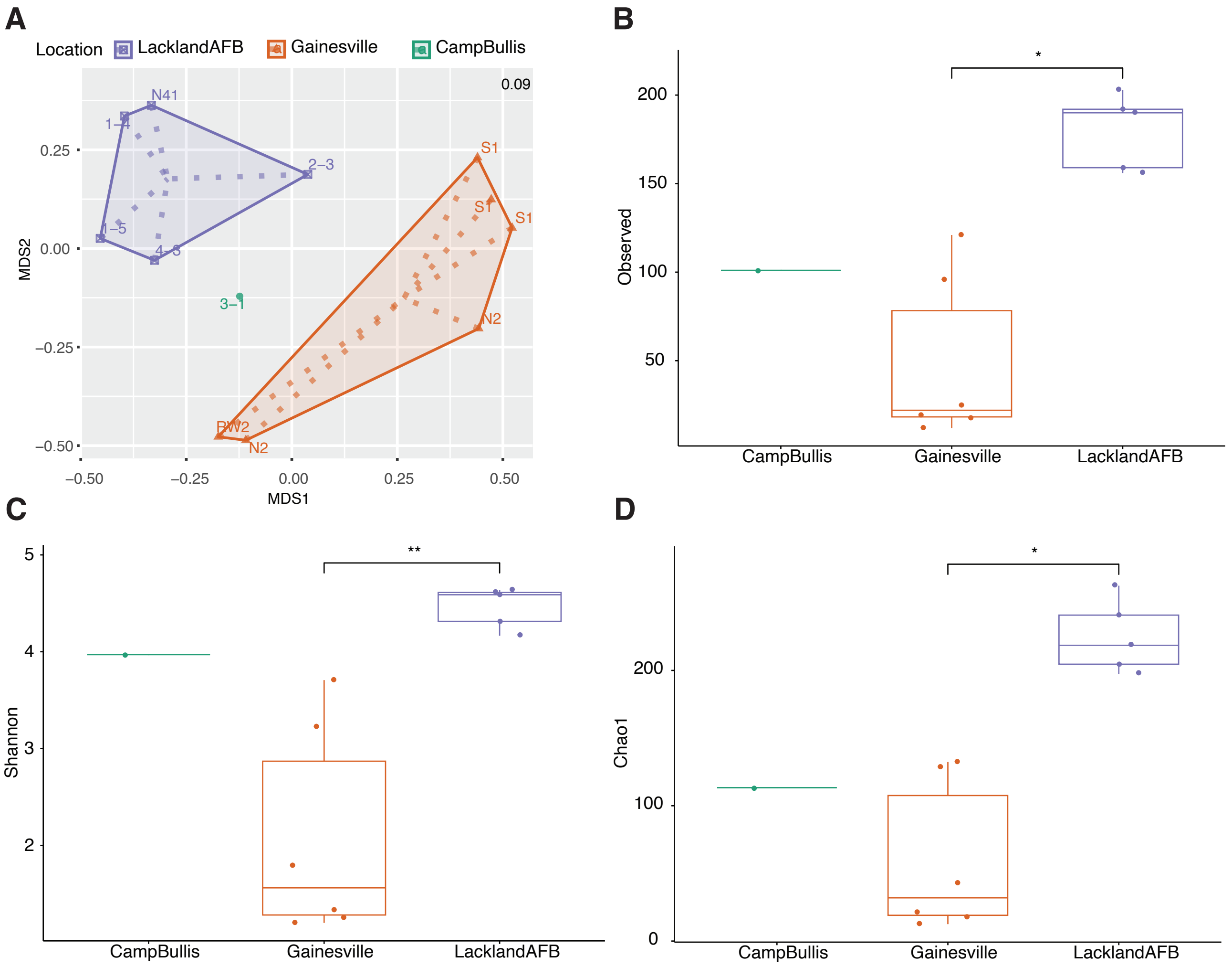


**Supplementary Figure S4:** Microbial profiles of nest materials. **A:** NMDS ordination for nest material samples based on Bray-Curtis dissimilarities. The number in the right upper corner indicates the stress value. Designation of each nest reflects the codes used in the heatmaps in Figure 1. **B:, C:, D:** Alpha diversity measures for nest material samples from different location. Asterisks stand for significant differences as identified by Dunn's Kruskal-Wallis multiple comparisons (Supplementary Table S1-DATASHEET S10)

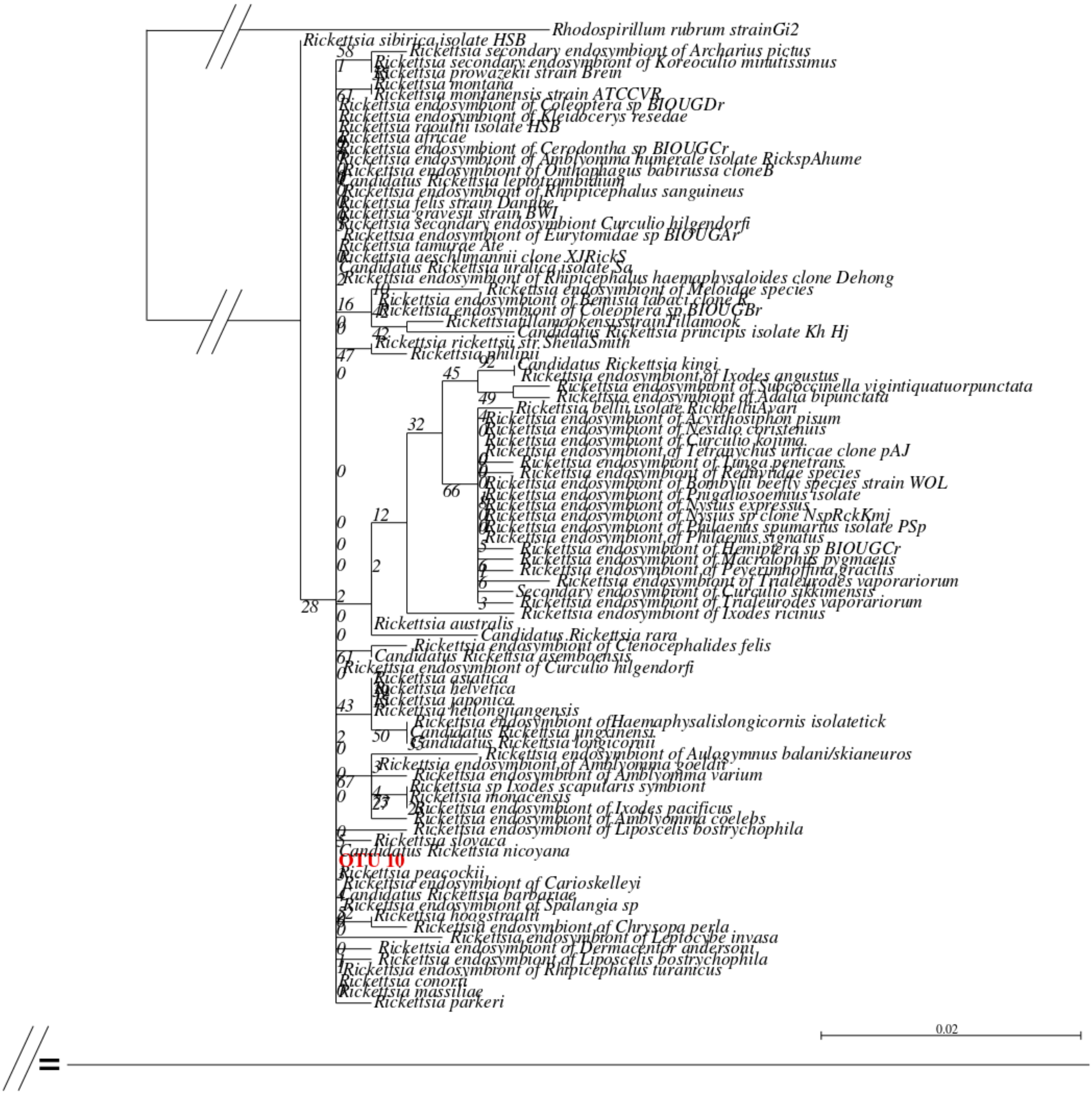


**Supplementary Figure S5:** Phylogenetic inference on 16S rRNA gene sequences of *Rickettsia* sp. detected in our samples. The sequences obtained for the 16S rRNA gene do not contain sufficient information to allow species assignment of OTU_10 (in red).

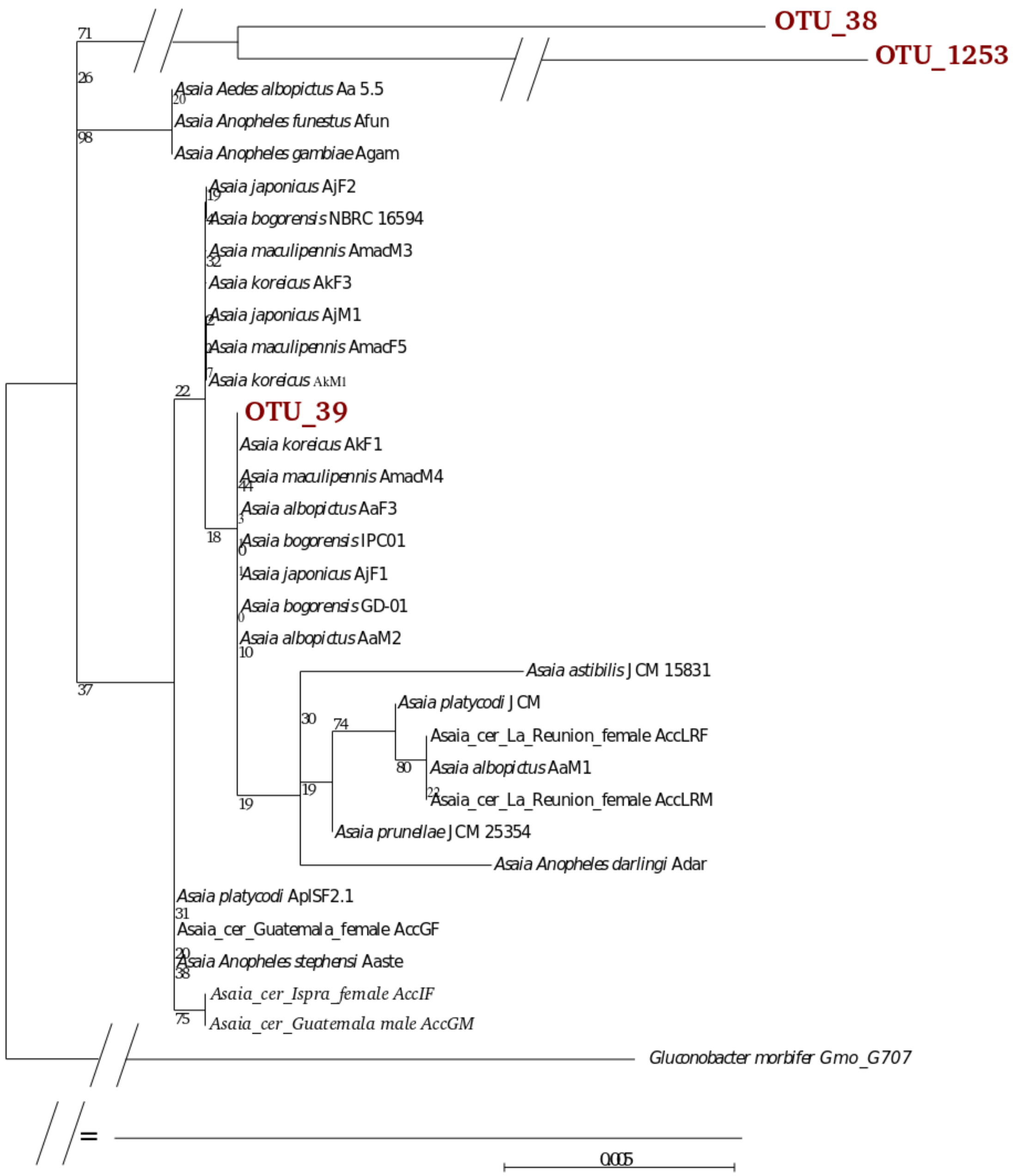


**Supplementary Figure S6:** Phylogenetic inference of *Asaia* OTUs based on 16s rRNA sequences. Amplicon sequences from our dataset are in red. OTU_39 clusters with *Asaia* strains detected in mosquitoes, whereas OTU_38 and OTU_1253 form long branches and their taxonomy could not be resolved.

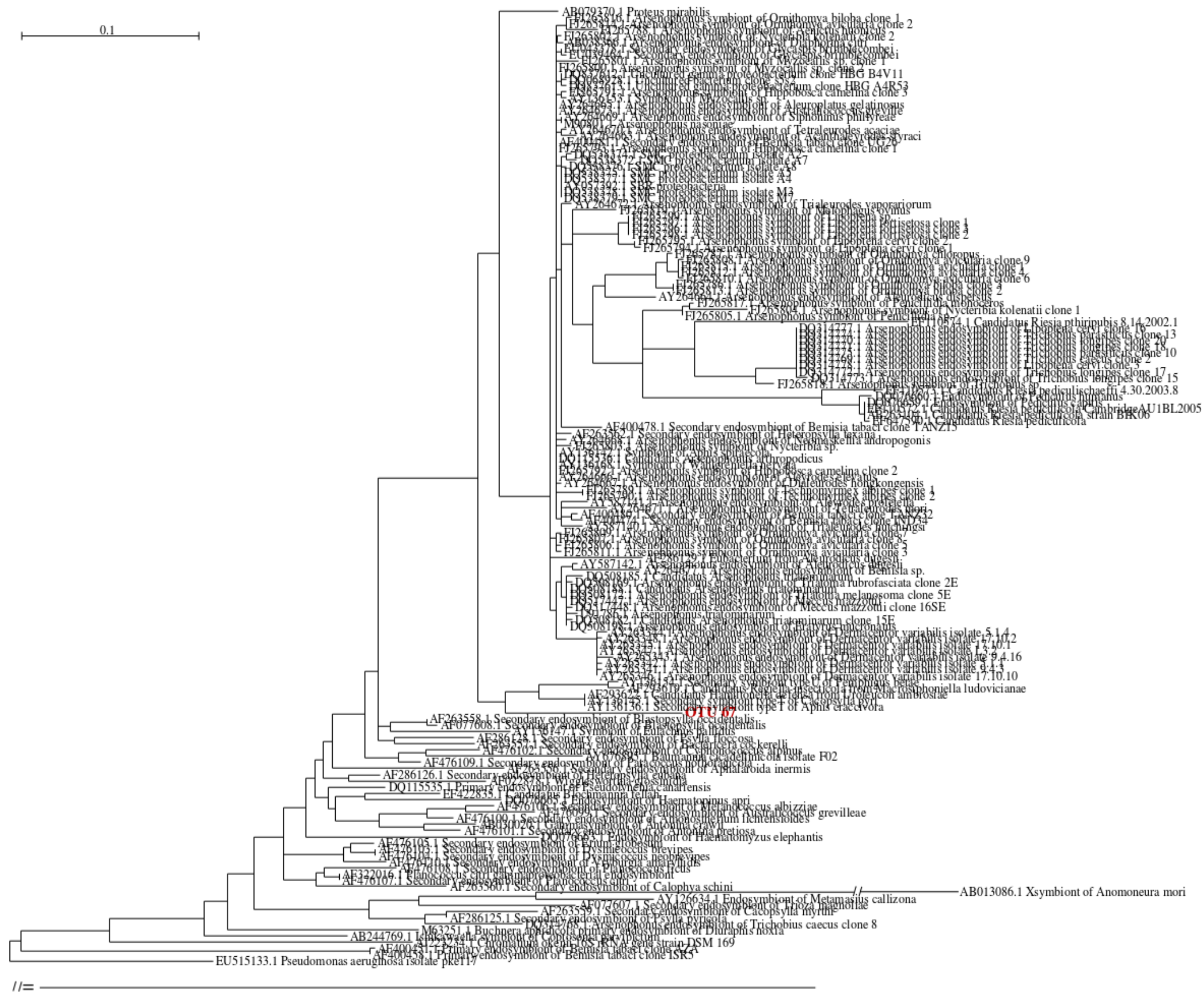


**Supplementary Figure S7:** Phylogenetic inference on 16S rRNA gene sequences of *Morganellaceae* detected in our samples. The sequences obtained for the 16S rRNA gene do not contain sufficient information to allow species assignment, although they place OTU_67 (in red) as sister group of symbionts of aphids.
